# CluStrat: a structure informed clustering strategy for population stratification

**DOI:** 10.1101/2020.01.15.908228

**Authors:** Aritra Bose, Myson C. Burch, Agniva Chowdhury, Peristera Paschou, Petros Drineas

**Affiliations:** Computational Genomics, IBM T.J. Watson Research Center, Yorktown Heights, NY, USA; Computer Science Department, Purdue University, West Lafayette IN, USA; Department of Statistics, Purdue University, West Lafayette IN, USA; Department of Biological Sciences, Purdue University, West Lafayette, IN, USA

**Keywords:** Population Structure, Association Studies, Clustering Ridge Regression

## Abstract

Genome-wide association studies (GWAS) have been extensively used to estimate the signed effects of trait-associated alleles. Recent independent studies failed to replicate the strong evidence of selection for height across Europe implying the shortcomings of standard population stratification correction approaches. Here, we present CluStrat, a stratification correction algorithm for complex population structure that leverages the linkage disequilibrium (LD)-induced distances between individuals. CluStrat performs agglomerative hierarchical clustering using the Mahalanobis distance and then applies sketching-based randomized ridge regression on the genotype data to obtain the association statistics. With the growing size of data, computing and storing the genome wide covariance matrix is a non-trivial task. We get around this overhead by computing the GRM directly using a connection between statistical leverage scores and the Mahalanobis distance. We test CluStrat on a large simulation study of discrete and admixed, arbitrarily-structured sub-populations identifying two to three-fold more true causal variants when compared to Principal Component (PC) based stratification correction methods while trading off for a slightly higher spurious associations. Applying CluStrat on WTCCC2 Parkinson’s disease (PD) data, we identified loci mapped to a host of genes associated with PD such as BACH2, MAP2, NR4A2, SLC11A1, UNC5C to name a few.

**Availability and Implementation:** CluStrat source code and user manual is available at: https://github.com/aritra90/CluStrat

## 1 Introduction

The basic principle underlying Genome Wide Association Studies (GWAS) is a test for association between genotyped variants for each individual and the trait of interest. GWAS have been extensively used to estimate the signed effects of trait-associated alleles, mapping genes to disorders and over the past decade about 10,000 strong associations between genetic variants and one (or more) complex traits have been reported [48,51,45,17]. One unambiguous conclusion from GWAS is that for almost any complex trait that has been studied so far, genetic variation is linked with many loci contributing to the polygenic nature of the traits. Hence, on average, the proportion of variance explained at the single marker is very small [45].

One of the key challenges in GWAS are confounding factors, such as population stratification, which can lead to spurious genotype-trait associations [37,39,33]. If a dataset consists of individuals from different ethnic groups, then the genotype data will be characterized by genome-wide *linkage disequilibrium (LD)* between variants. LD models the fact that alleles at different loci are correlated in individuals from the same ethnic group. Population structure causes genuine genetic signals in causal variants for a particular trait of interest to be mirrored in numerous non-causal loci because of LD [23], resulting in spurious associations. A related phenomenon, the so-called cryptic relatedness, is caused by individuals who are closely related and often grouped together by standard population structure correction strategies, and also poses a serious confounding problem [18]. Two popular approaches for stratification correction while building the *Genetic Relationship Matrix (GRM)* [2,41] involve (i) including the principal components of the genotypes as adjustment variables [37,38], and (ii) fitting a *Linear Mixed Model (LMM)* with an estimated kinship or GRM from the individual’s genotypes [51]. Recently, three independent studies [40,5,43] failed to replicate the previously reported signals of directional selection on height in European populations, as seen in the GIANT consortium (253,288 individuals [49]) in the independent and more recently UK Biobank cohort (500,000 individuals [8]). They further showed that the GIANT GWAS is confounded due to stratification along the north to south axis, where strong signals of selection were previously reported. These recent studies highlight the need for more sophisticated tools for correcting for population stratification.

Our work proposes a simple clustering-based approach to correct for stratification better than existing methods. This method takes into account the linkage disequilibrium while computing the distance between the individuals in a sample. Our approach, called CluStrat, performs *Agglomerative Hierarchical Clustering (AHC)* using a regularized Mahalanobis distance-based GRM, which captures the population-level covariance (LD) matrix for the available genotype data. We test CluStrat on large simulation studies of discrete and admixed, complex-structured populations of over 1,000 individuals genotyped on over one million genetics markers (Single Nucleotide Polymorphisms or SNPs for short) and we observe that our approach has the lowest number of spurious associations in our simulations. Our approach also identifies two to three-fold more rare variants at causal loci when compared to standard stratification correction strategies. Of independent interest is a simple, but not necessarily well-known, connection between the regularized Mahalanobis distance-based GRM that is used in our approach and the leverage and cross-leverage scores of the genotype matrix (see Methods and Appendix A for details).

## 2 Materials and Methods

**Notation** In the remainder of the paper we let matrix **X** ∈ ℝ^*m*×*n*^ denote the genotype matrix (e.g., the minor allele frequency (MAF) matrix on *m* samples genotyped on *n* SNPs). The matrix is appropriately normalized as is common in population genetics analyses to have zero mean and variance one (columnwise). The vector *y* ∈ ℝ^*m*^ represents the trait of interest and its *i*-th entry is set to one for cases and to zero for controls (for binary traits). We let **X**_*i*∗_ denote the *i*-th row of the matrix **X** as a row vector and **X**_∗*i*_ denote the *i*-th column of the matrix **X** as a column vector. We represent the top *k* left singular vectors of the matrix **X** by the matrix **U**_*k*_ ∈ ℝ^*m*×*k*^ and we will use the notation (**U**_*k*_)_*i*∗_ to denote the *i*-th row of **U**_*k*_ as a row vector.

### 2.1 CluStrat

CluStrat provides an LD based clustering framework to capture the population structure and tests for association within each cluster, as described in Algorithm 1.

**Algorithm 1.**
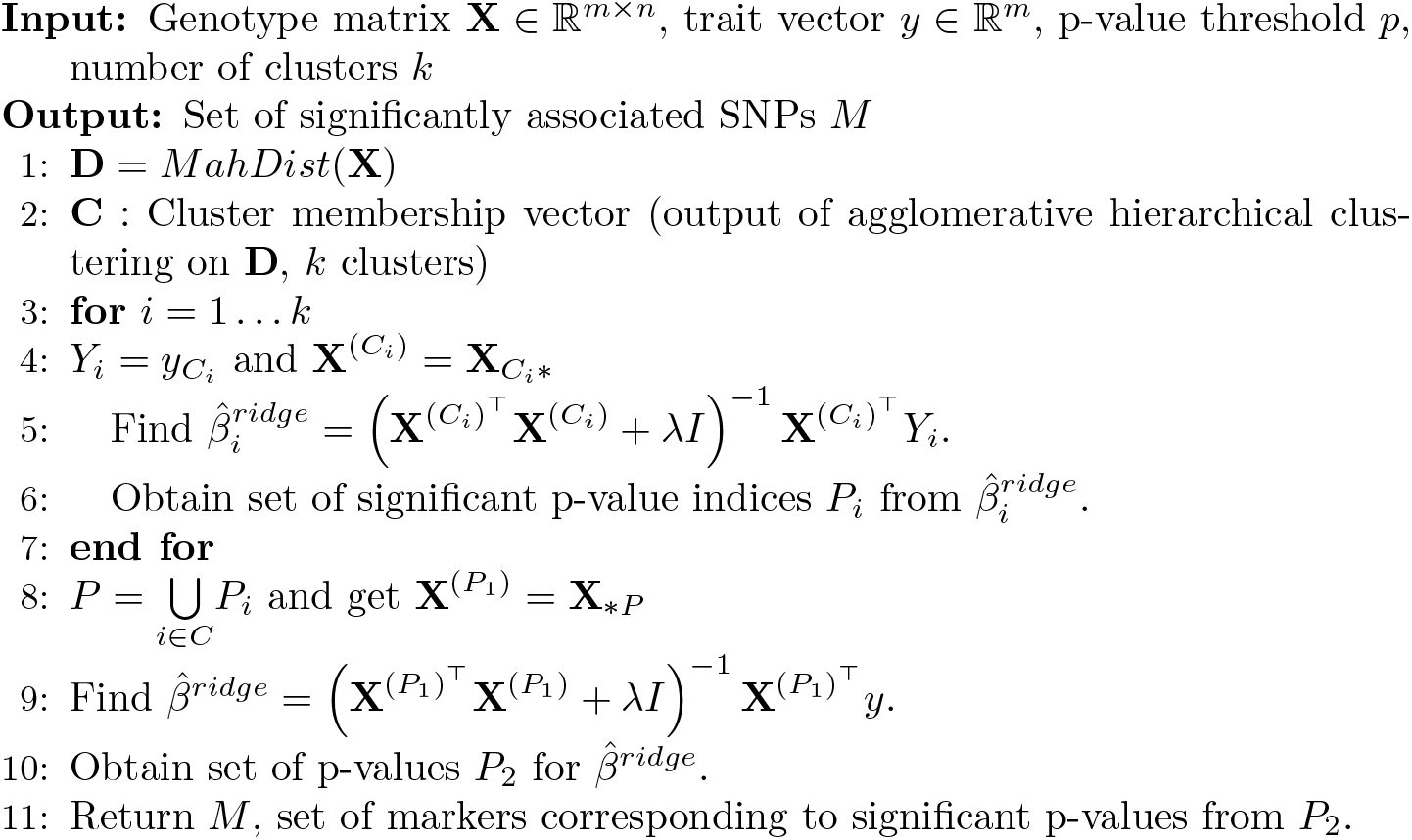
Structure informed clustering to correct for population stratification

It computes the distance matrix **D** from the normalized genotype matrix **X** and performs AHC for a number of clusters *k*, selected by a cross validation.

For each cluster, it runs an association test using ridge regression and obtains p-values for each marker. Thereafter, it computes *P*_1_ the union of intersections of significant associations across all clusters and select the corresponding markers from **X** to form 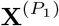. We can interpret this step as a scheme for variable selection. We run another association test with ridge regression on 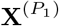 to obtain *M*, the final set of significant associations for all meta-analysis p-values below *p*.

We now briefly discuss the use of the Mahalanobis distance at the first step of the proposed algorithm. In an arbitrarily structured breeding population, correlation between loci due to LD often results in block-diagonal structures in the genetic relationship matrix. Thus, it is important to account for this LD structure in the computation of the distance matrix [34]. One way to account for the LD structure is to use the squared Mahalanobis distance [32,36] (denoted as **D** in eqn. 1). Given a matrix **G** ∈ ℝ^*n*×*n*^ which contains the covariance structure of LD (covariance due to LD between genetic markers), the LD-corrected GRM implementing the Mahalanobis distance is defined as

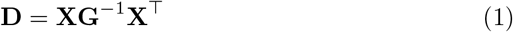

The Mahalanobis distance is useful in high-dimensional settings where the Euclidean distances fail to capture the true distances between observations (see Appendix A for relationships between Mahalanobis and Euclidean distances). It achieves this by taking the correlation structure between the features into account. We perform the association test in CluStrat by running ridge regression on each cluster. The regularizer, *λ*, is chosen by 5-fold cross validation. It is worth noting that we use ridge regression for each cluster as the number of samples is significantly smaller than the number of SNPs, thus making the overall system under-determined. We find the ridge-estimates as follows:

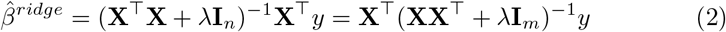

We emphasize that the above operation is run for each cluster. We simply dropped the superscripts from **X** in the above equation for simplicity. Then, we find the standard error of the estimates in order to calculate the p-values associated with each marker to compute the significance of its association with the trait. The standard error for each marker *i* in ridge regression is given by

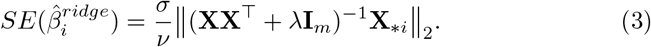

Recall that **X**_∗*i*_ is the *i*-th column of **X** and *ν* is known as the residual degrees of freedom. We set *ν* as shown in previous work [26] to the following,

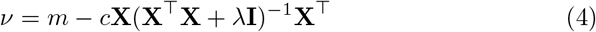

for a small scalar constant *c* > 0.

For biobank-scale datasets requiring terabytes of memory, computing the standard error can be a challenge. However, we can use random projection based approaches to sketch the input matrix **X** in order to approximate the standard error for each marker. This is indeed a novel contribution of our approach. We delegate details to Appendix A. We do note that our work is heavily based on previous work on Randomized Linear Algebra (RLA) [19,22,20,50]). To the best of our knowledge, this is the first approximation of the standard error in penalized regression using a sketching based framework and is of independent interest; see also [11] for related work.

### 2.2 Computing Mahalanobis Distance

Mahalanobis distance is known to be connected to statistical leverage [47]. We discuss the connection between a regularized version of the Mahalanobis distance and a regularized notion of statisical leverage scores below. We first note that the Mahalanobis distance is invariant to linear transformations, which means that the standard normalizations of the genotype matrix **X** do not affect the Mahalanobis distance between two vectors. Recall the definition of the Mahalanobis distance between samples *i* and *j*:

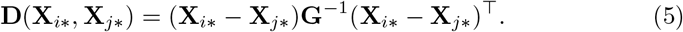

Now, recall that the rank-*k* leverage scores of the genotype matrix **X** ∈ ℝ^*m*×*n*^ with *n* ≫ *m* are defined by the row norms of the matrix of its top *k* left singular vectors **U**_*k*_ ∈ ℝ^*m*×*k*^. Let (**U**_*k*_)_*i∗*_ denote the *i*-th row of the matrix **U**_*k*_. Then the rank-*k* statistical leverage scores of the rows of **A**, for *i* ∈ 1, ···, *n* are given by

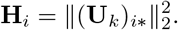

Similarly, the rank-*k* (*i*, *j*)-th cross-leverage score, **H**_*ij*_, is equal to the dot product of the *i*-th and *j*-th rows of **U**_*k*_, namely

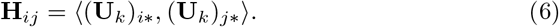

Here, **H** ∈ ℝ^*m*×*m*^ is the matrix of all leverage and cross-leverage scores. We note that 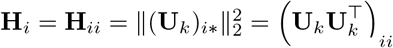 is a special case of the dot product in eqn. 6 for the diagonal leverage scores. We show that the Mahalanobis distance can be written in terms of the rank-*k* leverage and cross-leverage scores (see Appendix A for details on the relationship between Mahalanobis distance and leverage scores). Indeed, the final formulas are:

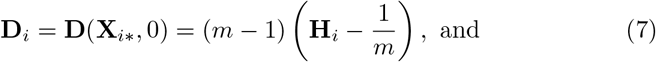

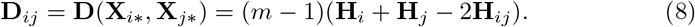

Thus, we show that Mahalanobis distance between two vectors can be computed efficiently without storing or inverting **G**, by the corresponding rank-*k* leverage and cross-leverage scores. By computing the rank-*k* Mahalanobis distance with respect to the top *k*-left singular vectors of the genotype matrix **X**, we make this computation feasible for UK Biobank-scale datasets using methods such as TeraPCA [7] to approximate the matrix **U**_*k*_ accurately and efficiently.

**Algorithm 2.**
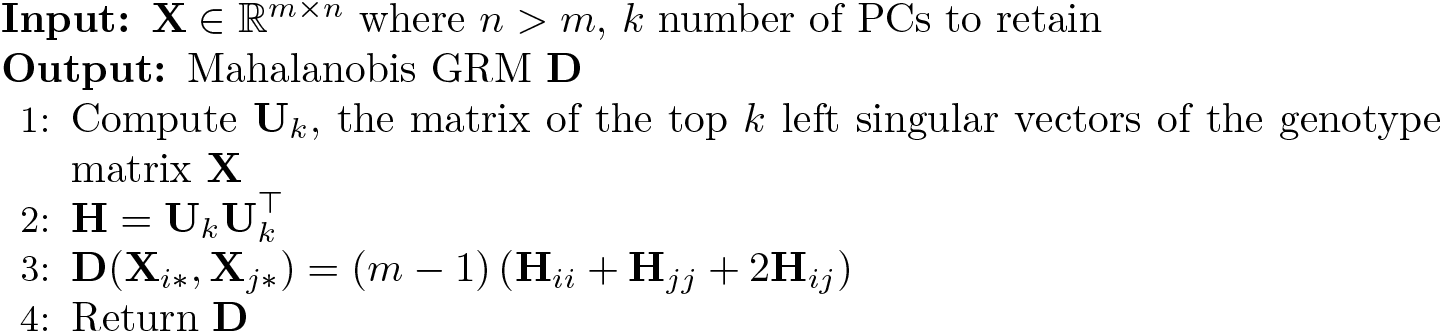
MahDist : Compute Mahalanobis distance based GRM

### 2.3 Agglomerative Hierarchical Clustering

We performed AHC using the LD induced Mahalanobis distance with a varying number of clusters. We set the expected number of clusters to *d*+*q* where *d* is the number of populations in the data and *q* is a user-defined range. We performed a five-fold crossvalidation to choose the optimal number of clusters and retain the cluster which maximizes the intersection of associations across all the clusters. The observed number of clusters is obtained by the inconsistency method of pruning according to the depth of the dendrogram. We note that for the simple case where *q* is set to zero, the clustering essentially attempts to recover the populations. In practice, we observed that the number of qualitative clusters obtained by running PCA on the genotype data serves as a good heuristic for the number of user defined clusters using the AHC procedure.

### 2.4 Datasets

We generated an extensive set of simulations with challenging scenarios to demonstrate the robustness to different real-world scenarios and power to detect few spurious associations.

We simulated and analyzed 100 GWAS datasets from a quantitative trait model (and it’s equivalent binary trait model using the Odds Ratio (OR) as the classifier for disease status from the continuous variable *y*) based on previous work [41].

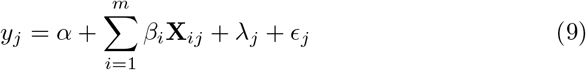

where *β*_*i*_ is the genetic effect of SNP *i* on the trait, *λ*_*j*_ is the random non-genetic effect and *ϵ*_*j*_ is the random noise variation for individual *j*. **X**_*ij*_ is the *i*^*th*^ marker for the *j*^*th*^ individual and *y* ∈ ℝ^*m*^ is the trait response variable (binary or continuous). For the genotype data, we simulated allele frequencies using (i) Balding-Nichols (BN) model [4] based on allele-frequency and *F*_*ST*_ estimates calculated on the HapMap data set, (ii) three different levels of admixture by varying the parameter *α* from {0.01,0.1,0.5} in Pritchard-Stephens-Donnelly model (PSD) [39] and (iii) structure estimated from 1000 Genomes Project (TGP) [3] (see Appendix A for details). To capture real world population structure, we applied CluStrat on the Parkinson’s Disease (PD) data available from the The Wellcome Trust Case Control Consortium (WTCCC2) study containing 4706 individuals (2837 controls and 1869 cases) across 517,672 SNPs. After performing quality control (details in Appendix) and pruning for LD between variants, we obtained 99,631 markers.

## 3 Results

We applied CluStrat to 30 simulation scenarios, modelling variable proportions of true genetic effect and admixture and compared its performance to standard population structure correction approaches such as EIGENSTRAT [38], GEMMA [52], and EMMAX [28]. We compared all the methods on all of the above scenarios with the p-value threshold set to 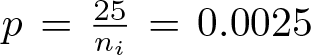; here *n*_*i*_ is the number of SNPs, which was set to 10,000. The expected number of spurious association as mentioned in [41] is *n*_0_ × *p* where *n*_0_ is equal to *n* minus number of causal SNPs (10 in our case). In all of the above scenarios, CluStrat outperformed the standard approaches in detecting the true causal variants while reporting slightly more spurious associations in the simulation scenarios.

The BN and PSD model simulates scenarios with unrelated isolated populations (Figure 2 (i) and (ii)). The samples when projected on the top two PCs clearly resemble three isolated clusters with no connections between them in BN and with admixed populations between the clusters in PSD, respectively. This serves as a “base case” for arbitrarily structured population with and without admixture. Armitage trend *χ*^2^ test with no population structure correction renders almost half of the SNPs in the simulation study as true associations, resulting in a vast number of spurious associations, clearly highlighting the need for population structure correction. PCA or LMM based approaches on the other hand return roughly the expected number of spurious associations as also shown in prior work [38]. Yet, PCA and LMM approaches are very stringent and detect zero causal SNPs in almost our experiments (Appendix figure 4, 6 and 7). CluStrat, however, strikes a balance between the two: it generates more spurious associations than the expected value, though far less than the Armitage trend *χ*^2^ test, and recovers almost similar number of causal SNPs. This shows that in an ideal case of population stratification correction, CluStrat can identify more causal SNPs mainly due to the use of the Mahalanobis distance and the simple clustering algorithm that we propose.

The TGP model is a more realistic model, drawing genotypes from allele frequency distributions from the 1000 Genomes Phase 3 dataset [3]. Projection of genotypes drawn from the 1000 Genomes (TGP) dataset on the top two axes of variations shows the distribution of samples across the world (see Figure 2 (iii). CluStrat (Figure 3) captures two-to-three fold increase in detection of causal variants while allowing for slightly more number of spurious associations. This shows that structure informed clustering of the genotype data followed by regularized association tests outperforms genotype and phenotype adjustments with the top *k* PCs, which is what EIGENSTRAT and LMMs often do.

Applying CluStrat on WTCCC2’s with a p-value threshold set to 10^−7^ we found a host of associated regions mapped to genes previously known to be associated with PD in literature. Our strongest associated loci rs10177996 (p-value = 2.2 × 10^−16^) maps to WNT10 in the Wignless-type MMTV integration site (Wnt) signalling cascade which has emerged as a very important pathway in major neurodegenerative pathologies including PD [29]. Another significant loci appeared to be in Chromosome 6 at rs176713 associated to the transcriptional inhibitor BACH2 which is known to interfere with Nrf2 function which when activated is a promising protective mechanism for progressive neurodegeneration in PD [27]. Other significant associated markers which are mapped to genes shown to be associated in previous work are SLC11A1 [6], UNC5C [30], MAP2 [16], EFNA5 [46], NR4A2, GRM7 [24], CNTN2, etc. (see Table 1 in Appendix B for details). We conclude that CluStrat not only works better in simulation scenarios but can also replicate previously recorded associations in real data sets such as PD.

**Table 1:**
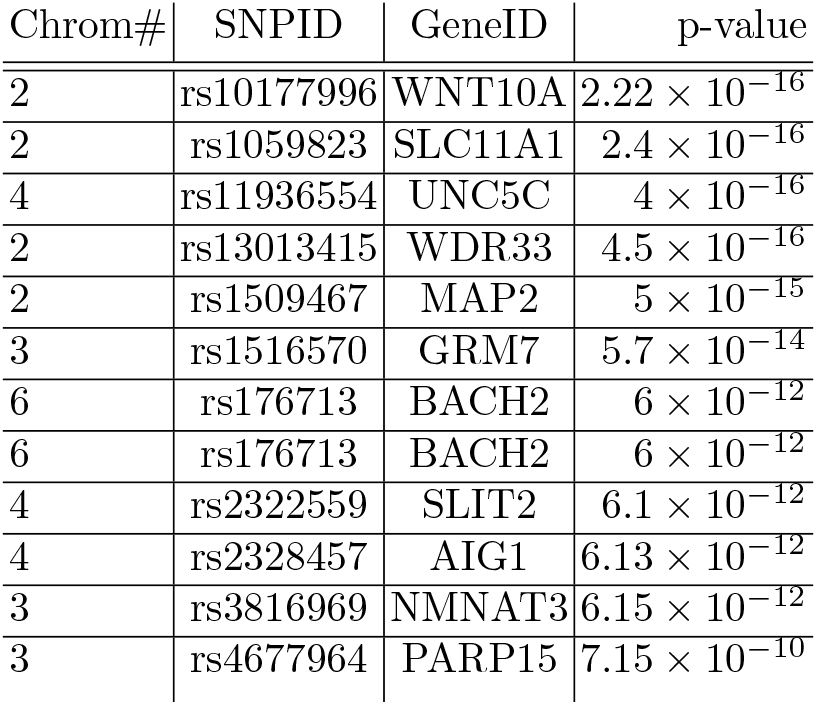
Table showing strongest associations after running CluStrat on WTCCC2 PD data

## 4 Discussion

CluStrat provides a structure informed clustering approach to correct for population stratification in GWAS. In our experiments, we verified the power of our approach in a variety of simulated data and observed that CluStrat outperforms the widely used EIGENSTRAT, GEMMA, and EMMAX methods in all settings, by detecting two to four times more causal SNPs. Adversely, our approach detects more spurious associations than standard approaches; however, it still performs much better than the uncorrected Armitage trend *χ*^2^ tests. Principal component based methods have been under scrutiny recently as independent studies [5,40] on the UK Biobank [8] failed to replicate the genetic associations of heritable height in Europeans, where a positive selection signal was observed in a north to south gradient [15,42,35] in the GIANT [49] cohort. These studies attributed the failure to replicate the results to cryptic relatedness among individuals, which PCA-based approaches for population stratification correction do not always capture. CluStrat provides a fine structure-based clustering approach to tackle cryptic relatedness and ancestral differences among the individuals between and within populations.

We chose to use the low-rank Mahalanobis distance metric in CluStrat because it captures the LD-induced structure information in the GRM. We established a link between the low-rank Mahalanobis distance and the low-rank leverage/cross-leverage scores, which allows us to get around the storage and computational bottlenecks of Mahalanobis distance. Prior work [34] computed the Mahalanobis distance by randomly subsampling a small number of SNPs to estimate the covariance matrix and circumvent to computational time and space requirements. Mahalanobis distance is also shown to remove bias in heritability estimates in presence of LD, therefore finding true causal variants [31]. An interesting topic for future work is to make CluStrat even faster by approximating the leverage and cross-leverage scores as shown in [19]. We also want to explore meta-analysis strategies to combine p-values from each cluster and obtain a cumulative significance across clusters as done in GWA studies. We showed that the Mahalanobis distance performs better (Figure 1) in capturing cryptic relatedness than the Euclidean distance based GRM. In upcoming work, we intend to evaluate CluStrat on the UK Biobank data to explore whether it succeeds or fails to replicate the north to south gradient of positive selection of height in Europeans [5,40]. CluStrat being a clustering based strategy is computationally slower than the PCA based approaches, we intend to explore Mahalanobis distance based GRM in other statistical models for association tests. Another future direction for CluStrat is to extend it to compute Polygenic Risk Scores (PRS) on a discovery or validation dataset and compare it with widely used packages such as PRSice2 [10] and LDPred [44], which compute PRS from GWAS summary statistics as well as raw genotypes.

**Fig. 1:**
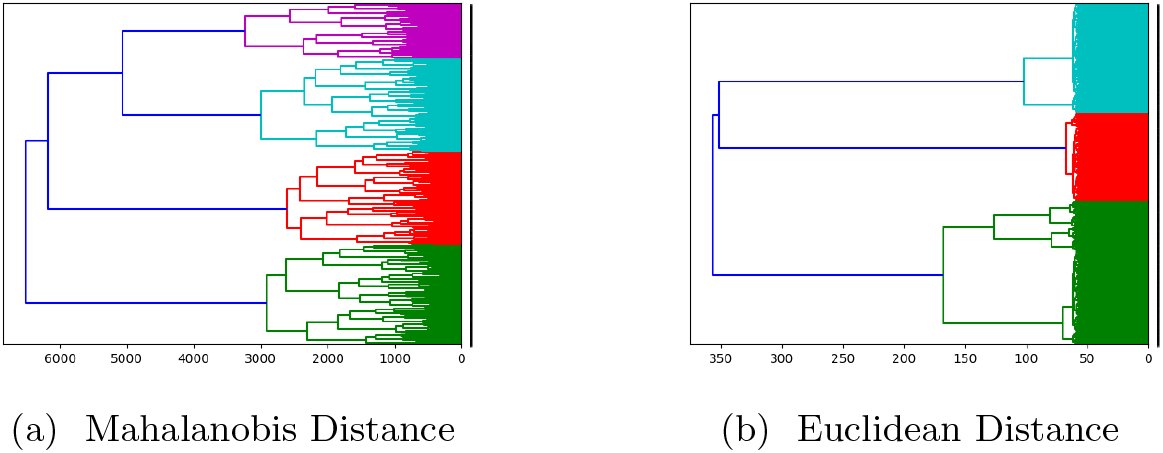
Dendrograms obtained after running AHC with Ward’s linkage on Pritchard-Stephens-Donnelly (PSD) model (*α* = {0.1, 0.1, 0.1}) shows Mahalanobis distance with fine grained interactions between the individuals inside a cluster recovering population substructure and cryptic relatedness which Euclidean distance based GRM fails to recover.

**Fig. 2:**
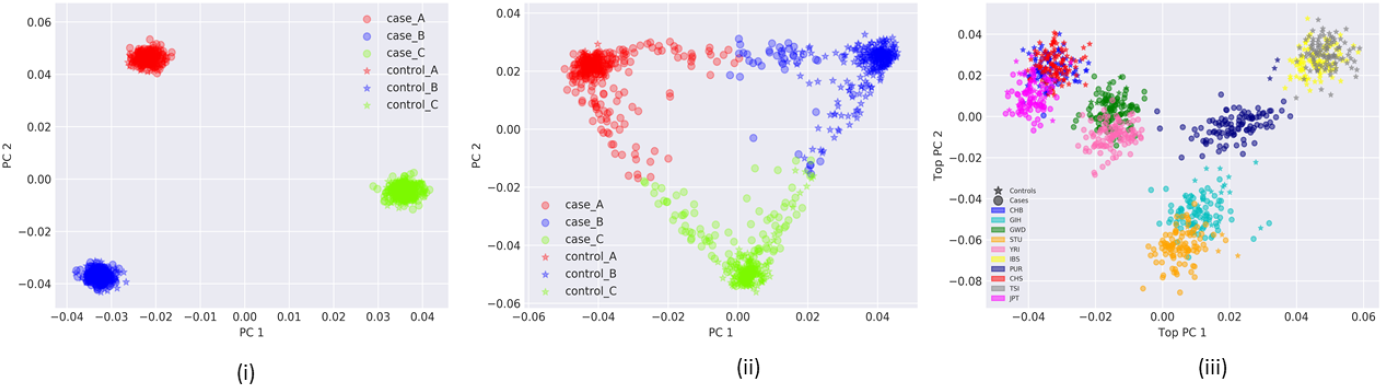
Projection of the samples from three populations simulated from (i) BN (ii) PSD (*α* = {0.1, 0.1, 0.1}) and (iii) TGP model on the top two axes of variation.

**Fig. 3:**
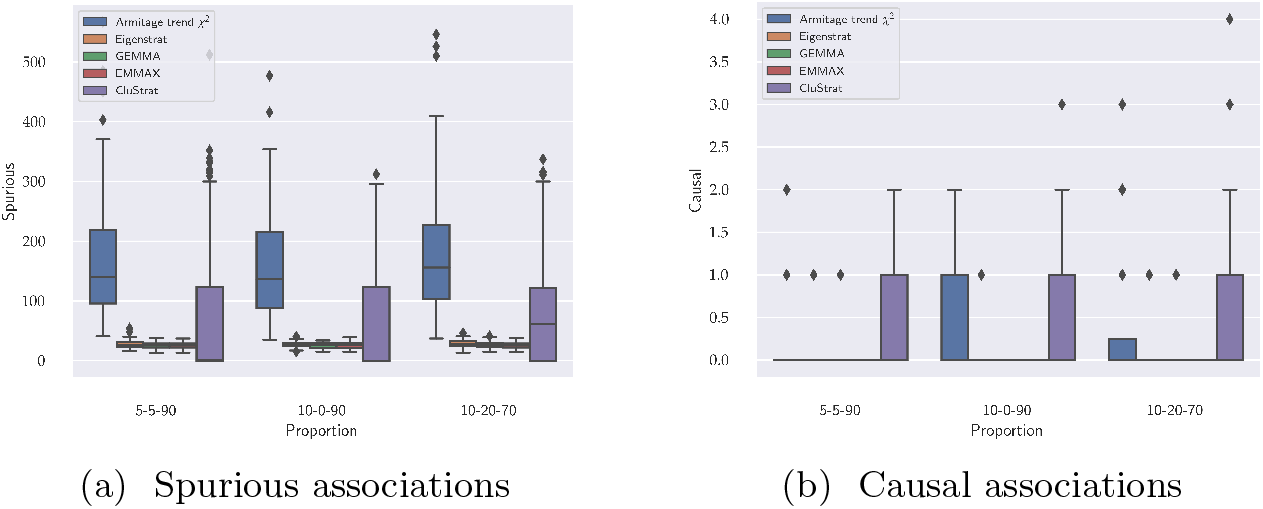
Box plots for spurious and causal associations on the TGP model shows Armitage trend *χ*^2^ has the maximum number of spurious associations (Appendix figure 4 and 6). CluStrat outperforms both the methods in this scenario by detecting two fold more causal loci while allowing slightly more spurious associations.

**Fig. 4:**
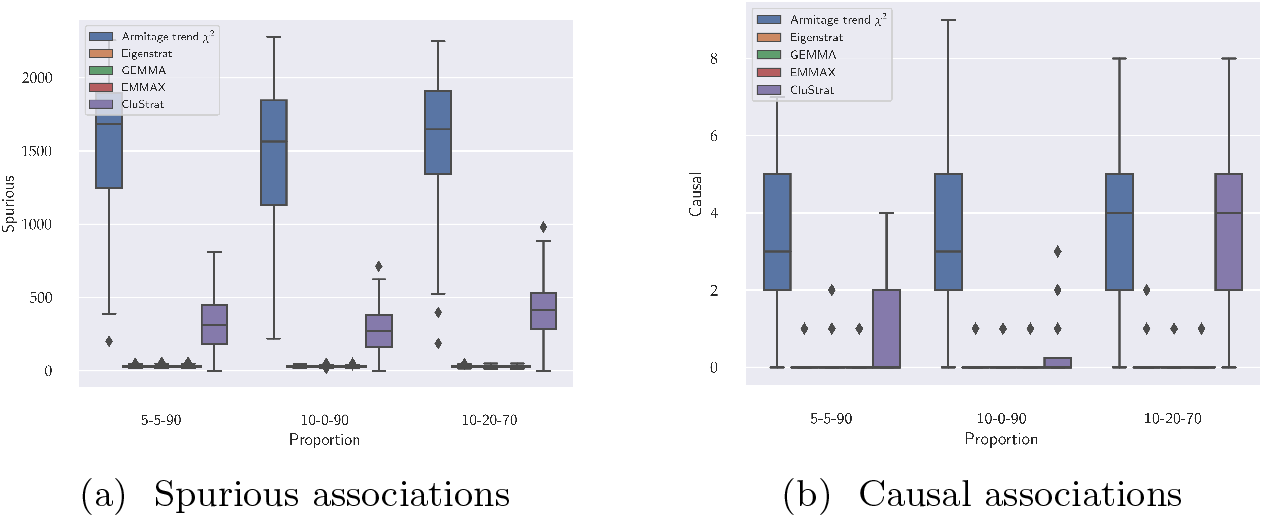
Box plots for spurious and causal associations on the BN model shows that Armitage trend *χ*^2^ has the maximum number of spurious associations containing about 4-5 causal SNPs whereas EIGENSTRAT has minimum number of spurious associations while detecting almost zero causal SNPs. CluStrat has more spurious associations than EIGENSTRAT and considerably less than Armitage trend *χ*^2^ recovering slightly more number of causal SNPs than the latter.

In summary, CluStrat highlights the advantages of biologically relevant distance metrics, such as the Mahalanobis distance, which seems to capture the cryptic interactions within populations in the presence of LD better than the Euclidean distance. We evaluated CluStrat on a host of simulation scenarios for arbitrarily structured populations with and without admixture. The choice of the number of clusters does not change the results drastically and one can use the number of broad clusters that are visually apparent when plotting the data on the top two or three principal components as a initial choice of clusters. We implemented a five-fold cross validation approach to obtain the optimal choice for the number of clusters and the regularization parameters. We concluded that CluStrat outperforms PCA or LMM based population stratification correction techniques in a variety of simulated datasets.

## 5 Acknowledgements

AB carried out this work as a part of his PhD dissertation in the Computer Science Department, Purdue University, West Lafayette, IN, USA.

## APPENDIX A Data simulator

The complete simulation study on quantitative traits with population structure latent variable is constructed in 5 different ways for 3 different proportions of variance among genetic effects, non-genetic effects and random noise, all of which contributing to the trait. We simulated 100 independent datasets containing *m* = 1, 000 individuals and *n* = 100, 000 markers from a quantitative trait model 9. Let *Z* be a latent variable which captures environmental factors contributed by population structure. Equation 9 allows interdependence of structure, lifestyle and environment. We assume 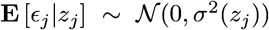 allowing for heteroskedasticity of the random noise variation [41]. Therefore, 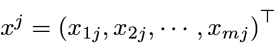, *λ*_*j*_ and *σ*^2^ can be thought of as functions of *z*_*j*_ where 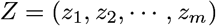. *λ*_*j*_ is unspecified but along with *z*_*j*_, they are assumed to be dependent, random variables. Thus, the population genetic model is dependent on the structure variable *z*_*j*_ for each individual. We define the corresponding binary trait model as

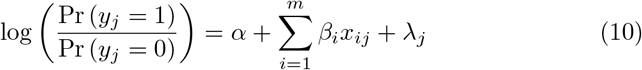

using the Odds Ratio (OR) as the classifier for disease status from the continuous variable *y*.

The complete simulation study on quantitative traits with population structure latent variable is constructed in 5 different ways for 3 different proportions of variance among genetic effects, non-genetic environmental effects and random noise, al of which contributing to the trait. Therefore 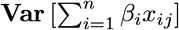, 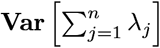 and **Var**[*ϵ*_*j*_] are assigned in proportions of (5%,5%,90%), (10%,0%,90%) and (10%,20%,70%), respectively. Thus, we varied the amount of genetic contribution to the trait for each simulation scenarios and capture variable amounts of population structure confounding. We simulated ten truly associated SNPs whose effect sizes were distributed according to a Normal distribution and we set *β*_*i*_ = 0 for all other non-causal SNPs.

The genotype matrix **X** ∊ ℝ^*m*×*n*^ consisting of the simulated allele frequencies was simulated using the algorithm from a previous study [25,41]. Specifically, we set **F** = **TS** where **T** ∊ ℝ^*m*×*d*^ and **S** ∊ ℝ^*d*×*n*^ where *d* ≤ *n* is the number of population groups. **S** is the matrix containing the population groups encompassing the structure for the individuals shared across all SNPs. On the other hand, **T** characterizes how the structure is manifested in the allele frequencies of each SNP [25]. Finally, projecting **S** onto the column space of **T** we obtain the allele frequency matrix **F**. We sample **X** as a special case of **F** for Balding-Nichols (BN), Pritchard-Stephens-Donelly (PSD) and TGP (1000 Genomes Project), respectively. We formed **T** and **S** for the above 5 simulations with 3 scenarios each and continuous traits, resulting in, 15 different evaluation scenarios each for continuous and binary traits. The algorithm for constructing **T** and **S** is detailed in reference [25,41].

For BN, the allele frequency matrix is simulated from the HapMap phase 3 dataset [14] using three unrelated populations. The final genotype matrix, **X**, is drawn independently at random from the Binomial distribution with parameters *n* set to 2, denoting the allele status (0,1 or 2) corresponding to homozygous major/minor or heterozygous with probability *p* set to the simulated allele frequency for each individual-SNP pair. For PSD, the allele frequency matrix was drawn from the BN frequency distribution. However, it differs from BN in simulating **S** by i.i.d draws from Dirichlet distribution with varying *α* which denotes the parameter influencing the relatedness between the individuals. We show results for *α* = {0.01, 0.1, 0.5} here and conducted simulations on a wide range of *α* values from 0.01 to 0.5.

## Real dataset

To capture real world population structure, we applied CluStrat on the Parkinson’s Disease (PD) data available from the The Wellcome Trust Case Control Consortium (WTCCC2) study containing 4706 individuals (2837 controls and 1869 cases) across 517,672 SNPs. After performing quality control by filtering for genotyping rate lower than 99%, MAF less than 0.01 and Hardy-Weinberg equilibrium less than 0.001 and pruning for LD between variants higher than 0.2 squared correlation we obtained 99,631 markers.

## Distance metrics for Hierarchical clustering

CluStrat computes the distance matrix **D** from **X** to perform the AHC. The choice of distance metric is user defined. However, we choose the distance metric based on LD induced distances to capture the cryptic relatedness between individuals in a population which is not otherwise captured by other stratification methods. We use the normalized genotype matrix **X** following the standard normalization procedure by minor allele frequency of each marker. Let us consider the unscaled GRM which captures the Euclidean distances as **D** = **XX**^⊤^ and let **I** ∈ ℝ^*n*×*n*^ (in the order of the number of markers *n*). Thus **D** can be rewritten as

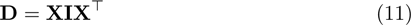

Thus we can see the unscaled GRM as the same weighting on the diagonal for all markers. In an arbitrarily structured breeding population, there exists correlation between loci due to linkage resulting in varying values along the diagonal or a block-diagonal structure in the GRM. Thus, it is important to account for this LD covariance structure in the computation of the GRM [34]. One way to account for the LD structure in GRM is to use the squared Mahalanobis distance [32,36] (denoted as **D** for simplification). Given a matrix **G** ∈ ℝ^*n*×*n*^ which contains the covariance structure of LD (covariance due to markers), then the LD-corrected GRM with Mahalanobis distance is defined as

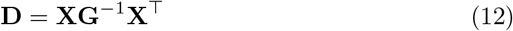

The RHS of equation 1 represents the squared multivariate Mahalanobis distance between individuals. Mahalanobis distance is useful in a high-dimensional setting where the Euclidean distances fail to capture the true distances between observations. It achieves this by taking correlation between the features captured in the SNP covariance matrix into account. The Cholesky factorization of the covariance matrix **G** = **LL**^⊤^ where **L** is the lower diagonal matrix known as the Cholesky factor of **G** [34]. We can represent equation 1 as

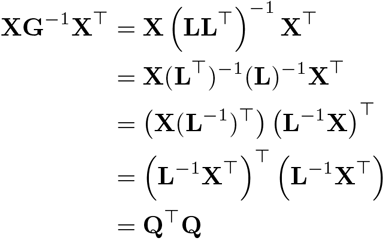

**Q** = **L**^−1^**X**^⊤^ represents the transformed variables and **Q**^⊤^**Q** is the squared Euclidean distance between the transformed variables. Thus, Mahalanobis distance accounts for covariance between variables by transforming the data into an uncorrelated form and computing the euclidean distances between them.

## Mahalanobis Distance and Leverage Scores

Mahalanobis distance is known to be connected to statistical leverage [47], which is extended in the RandNLA framework as leverage scores. We show this relationship by first noting that Mahalanobis distance is invariant to linear transformations, which means the Mahalanobis distance between two vectors,

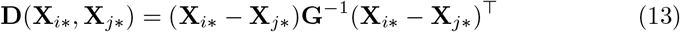

can have zero means for each vector. In our genotype matrix, **X** ∈ ℝ^*m*×*n*^, we have *n* markers and *m* observations. The design matrix **X** on which we intend to fit the model, however, must contain an intercept and thus we refer to **X** here as the design matrix containing the intercept column followed by one column for each SNP for all the individuals in rows. Furthermore, as we compute the Mahalanobis distance with respect to the *low-rank* genotype matrix **X**_*k*_, we only consider the *low-rank leverage scores* (rather than the leverage scores of the original matrix **X**) which are essentially the diagonal elements of the following projection-matrix:

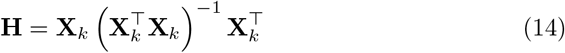

and similarly, the off-diagonal elements of **H** are called *cross-leverage scores* of **X**_*k*_. Now, we will give a clean connection between Mahalanobis distance and these leverage and cross-leverage scores.

First, consider the diagonal elements of **H** *i.e.* when *i* = *j*, we have

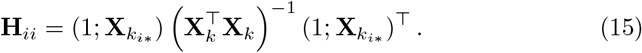

Exploiting the structure of 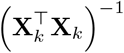, we can reformulate it in terms of a block matrix as follows

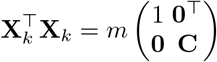

where 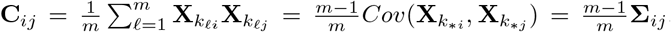. **Σ** here is the corresponding sample covariance matrix. Thus,

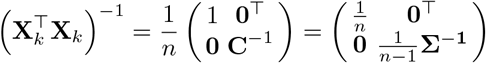

From Equation 15 we obtain

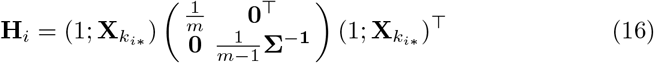

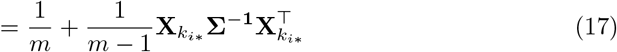

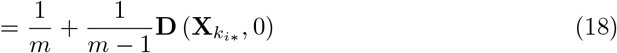

Solving for

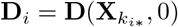

yields,

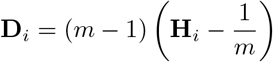

Similarly, we can prove the cross-leverage scores

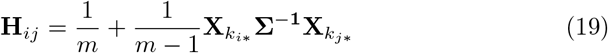

To prove the relationship of **H**_*ij*_ with **D**_*ij*_ we see,

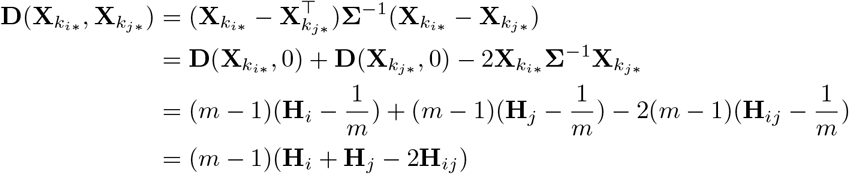

If we take 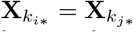 then we find 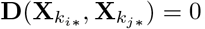. Thus, we show that Mahalanobis distance between two vectors can be computed by the corresponding vector’s leverage scores.

One of the key computational bottlenecks of Mahalanobis distance is computing the inverse of the SNP covariance matrix **G** as required in Equation 1. In real datasets, with the improvements in genotyping and sequencing technologies, the number of SNPs can be in the millions, thereby making **G** in the order of million times million and infeasible to store in secondary memory. Here, we propose the first approximation of Mahalanobis distance by computing leverage and cross-leverage scores in a faster and efficient way. As we have shown in Equation 19 and 16 following up from previous work [47], Mahalanobis distance can be written in terms of leverage scores. Advances in RandNLA community have brought about faster computations for leverage scores as well as cross-leverage scores; hence, we can compute approximations to these scores using random sampling algorithms with theoretical guarantees [19]. For our purposes of demonstrating the proof-of-concept, we work with simulated data as described above for 1,000 individuals and 500,000 SNPs which could be feasibly processed in a personal workstation to compute the deterministic leverage and cross-leverage scores.

## Computing leverage and cross-leverage scores

In fact, we do not need to compute the *rank-k* leverage and cross-leverage scores exactly. Using the idea of [19,12], they can be well-approximated in a much faster way with high-probability. In particular, computing *m* row-leverage scores takes time

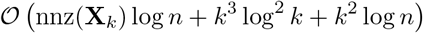

where nnz means the non-zero entries of the matrix, and computation of the high-valued cross-leverage scores can be done in time

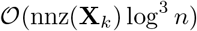

## Fast Computation of Standard errors

For biobank-scale data-sets requiring terabytes of memory, computing the standard error can be a challenge. However, we can use random projection based sketching matrices to find an approximate standard error for each marker by projecting the genotype matrix **X** on a sketching matrix **S** ∈ ℝ^*n*×*r*^ to form a sketch **XS**. We can rewrite the standard error in Equation 3 to find it’s approximate as,

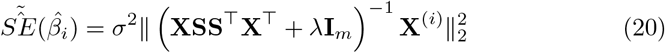

The sketched matrix **XS** generically has the same rank but much fewer columns than **X**, satisfying 1 ≤ *r* ≤ *min*{*m, n*}. Sketching, in general, is used to speed up solving systems of linear equations [20,22,12]. The sketching dimension, *r*, is directly proportional to the accuracy obtained by the approximate standard errors. Some prior knowledge of the design matrix, **X**, helps determine the target rank, *r*, that will result in satisfactory error guarantees. The sketching matrix, **S**, can be chosen simply as i.i.d normal random variables with mean equal to zero and variance equal to 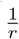. There exists other ways to choose **S** based on random projections as shown in previous work involving Fast Johnson-Lindenstrauss Transform [1], Subsampled Randomized Hadamard Transform [20,21] and Count-Sketch matrices [9] from streaming setting involving faster computation with sparse matrices.

## Time to compute eqn. (20)

Following the discussion as in [11], let the time to compute the sketch **XS** ∈ ℝ^*m*×*s*^ be *T* (**X**, **S**) which depends on the particular construction of **S**. In order to invert the matrix **Q** = **XSS**^⊤^**X**^⊤^, it suffices to compute the SVD of the matrix **XS**. Notice that given the singular values of **XS**, we can compute the singular values of **Q** and also notice that the left and right singular vectors of **Q** are the same as the left singular vectors of **XS**. Interestingly, we do not need to compute **Q**^−1^. Instead, we can store it implicitly by storing the left (and right) singular vectors of **Q** along with its singular values, **Σ**_**Q**_. Then, we can compute all necessary matrix-vector products using this implicit representation of **Q**^−1^. Thus, inverting **Q** takes 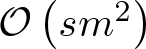 time and this will eventually dominate the computation of all other matrix-vector products and the Euclidean-norm. Therefore, total running time to compute eqn. (20) is given by

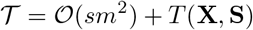

Clearly, specific constructions of the sketching matrix **S** will determine both *s* and *T*(**X**, **S**), and therefore 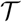. For example, if **S** is a subsampled randomized Hadamard transform (SRHT) matrix, then we have, 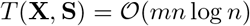 and 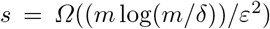; therefore 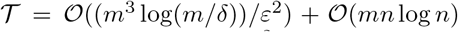. Similarly, if **S** has sub-Gaussian entries, then 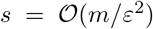 and 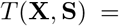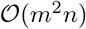; therefore 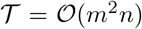. Furthermore, if **S** is a count-sketch matrix of [12], then, in this case, 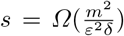 and *T*(**X**, **S**) = nnz(**X**). So, total running time 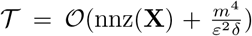. Here, *∊* is the accuracy parameter and *δ* is the corresponding failure probability.

Note that sketching-dimension, *s*, for sub-Gaussian is optimal, but *T*(**X**, **S**) takes much time. On the other hand, for count-sketch, *T* (**X**, **S**) is much faster (only nnz(**X**)), but sketching-dimension is huge 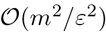. In a recent work [13], the authors showed that we can actually use all the sketches discussed here in conjunction with each other to get the best performance both in terms of sketching-dimension as well as computation time. More precisely, if one set **S** = **S**_1_**S**_2_**S**_3_ with **S**_1_ being the count-sketch, **S**_2_ being the SRHT and **S**_3_ being the sub-Gaussian, the **XS** will have 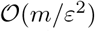 columns with running time 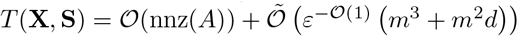.

## APPENDIX B Comparing Stratification Methods

## BN model

The BN model simulates scenarios with unrelated isolated populations (Figure 2 (i)) and serves as the basic case for arbitrarily structured population with no admixture.

The samples when projected on the top two PCs clearly resembles three isolated clusters with no connections between them. This is an ideal case when the populations are not mixing due to environmental factors acting as barriers of gene flow between populations. GWAS has shown to be robust in these settings [45]; however, the cryptic relatedness for each cluster remains a plaguing issue [5]. We ran CluStrat on this scenario with p-value threshold set to 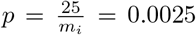 (*m*_*i*_ is the number of SNPs in each iteration, set to 10,000 for 100 iterations). The expected number of spurious association as mentioned in [41] is *m*_0_ × *p* where *m*_0_ = *m*− number of causal SNPs. In our case, as we set the number of causal SNPs to 10 as per [41], *m*_0_ = 9990 and therefore, the number of spurious associations to be approximately 25 with degree of freedom set to 1 for genotypes.

Armitage trend *χ*^2^ with no population structure correction renders almost half of the SNPs in the simulation study as true associations resulting in a considerable amount of spurious associations highlighting the need for population structure correction. EIGENSTRAT on the other hand results in the expected number of spurious associations as also shown in previous work [38]. However, it behaves stringently and detects zero causal SNPs almost all of the time (Figure 4). CluStrat, however, strikes a balance between the two and generates far more spurious associations than the expected value but about 5 folds less than Armitage trend *χ*^2^ recovering a slightly higher number of causal SNPs. This shows that in the ideal case of population structure correction, CluStrat can identify more causal SNPs due to the structure informed clustering setup which widely used stratification correction methods lack.

## PSD model

The PSD model emulates real world datasets more closely than BN model. It allows for admixing individuals and gradients across the populations. It is sampled from the Dirichlet distribution parameterized by a concentration parameter *α* ∈ ℝ^*d*^ where *d* = 3 (the number of populations for all simulations conducted). A higher value of *α*_*i*_ corresponds to greater weight of *i*^*th*^ population. We ran CluStrat on the PSD model with varying number of *α* from 0.01 to 0.5 and kept equal *α*_*i*_ for a symmetric distribution. We report the boxplots of spurious and causal associations (Figure 6 and 7) for *α* = 0.1, 0.5 and and observe that for the first case of variance, (5%, 5%, 90%), Armitage trend *χ*^2^ and CluStrat performs almost similarly in terms of spurious associations. This is due to the fact that only 5% of the trait is explained by true genetic associations in presence of LD and the rest is noise and environmental factors. However, CluStrat outnumbers EIGENSTRAT, GEMMA and EMMAX in terms of causal associations and detects four to six fold more true causal SNPs. For the other two variance proportions, CluStrat performed better than the other methods in detecting the causal associations and strikes a balance in terms of spurious associations.

## Comparing Distance metrics

CluStrat with Euclidean distance metric based GRM (sample covariance matrix) also contains structure information as part of the relationships between the individuals within and between population groups. The GRM with Euclidean distance is straightforward to compute as shown below

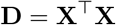

where **X** ∈ ℝ^*n*×*m*^, with number of markers, *n* and number of samples, *m* (*n* >> *m*). We show that although Euclidean distances between individuals is straightforward to compute, it fails to distinguish fine-grained distances between individuals in the same cluster owing to cryptic relatedness. This is highlighted after performing AHC using Ward’s linkage method which minimizes the increase in sum of squares between two cluster centroids in order to decide when to merge them. (Figure 1).

When Mahalanobis distance based GRM is used instead of Euclidean distance in AHC on PSD model with 1,000 individuals and 10,000 SNPs across 3 admixed arbitrarily structured ethnic groups, it reveals four broad clusters with various fine-grained sub-clusters revealing how Mahalanobis distance help recover cryptic relatedness and substructure within a population.

Due to admixture in the PSD model (*α* = {0.1, 0.1, 0.1}) as shown in Figure 5 the dendrogram finds three broad clusters owing to the three populations in the simulation. It subsequently finds different sub-clusters at different depth on the horizontal axis. Thus, identifying interaction between individuals inside a cluster. This is a significant advantage of using Mahalanobis distance over it’s Euclidean counterpart as the latter only reveals three broad clusters with indistinguishable interactions in each cluster (Figure 1).

**Fig. 5:**
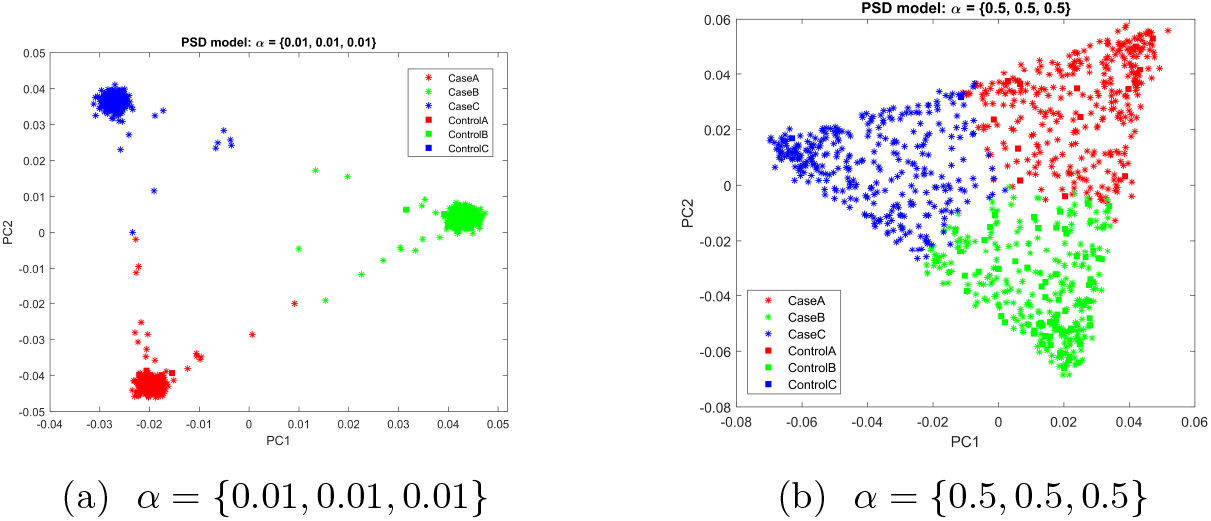
Projection of the samples from PSD model with varying sets of values of *α*. We observe that increasing *α* increases the density between individuals leading to admixture and creates a uniform gradient as all values of *α*_*i*_ are equal.

**Fig. 6:**
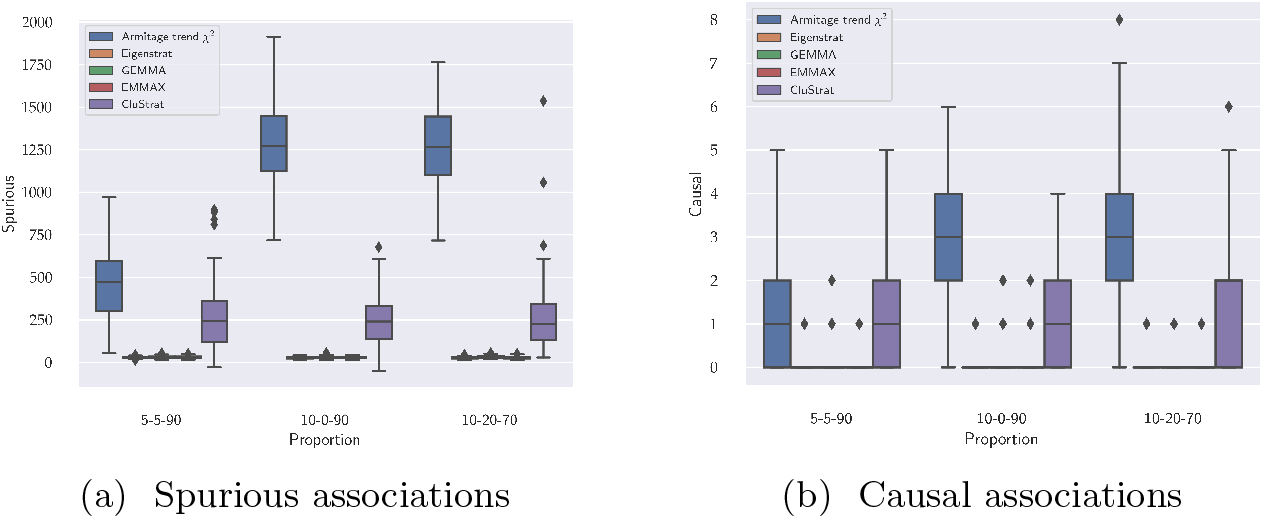
Box plots for spurious and causal associations on the PSD model (*α* = {0.1, 0.1, 0.1}) shows Armitage trend *χ*^2^ has maximum number of spurious associations containing less causal SNPs than the BN model (Figure 4) owing to the admixed nature of the individuals in PSD. EIGENSTRAT, GEMMA and EMMAX has least number of spurious associations while detecting almost zero causal SNPs. CluStrat has more spurious associations than the standard approaches and less than Armitage trend *χ*^2^ while recovering two to three fold more causal SNPs.

**Fig. 7:**
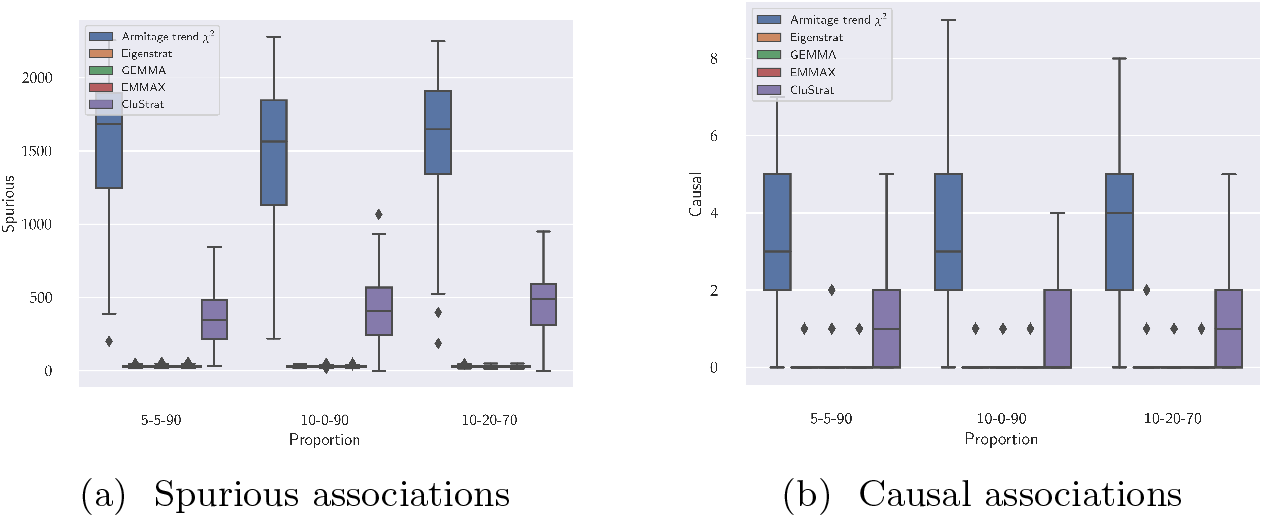
Box plots for spurious and causal associations on the PSD model (*α* = {0.5, 0.5, 0.5}) shows Armitage trend *χ*^2^ has maximum number of spurious associations containing less causal SNPs than the BN model (Figure 4) owing to the overtly admixed nature of the individuals in PSD. EIGENSTRAT, GEMMA and EMMAX has least number of spurious associations while detecting almost zero causal SNPs. CluStrat has more spurious associations than the standard approaches and slightly more than *α* = 0.1 owing to more admixed nature of the data. It has considerably less spurious associations than Armitage trend *χ*^2^ while recovering two to three fold more causal SNPs.

When we ran AHC with both the distances, we observe similar performance on the PSD model with Mahalanobis distance based GRM performing slightly better with respect to it’s Euclidean counterpart (Figure 8). We note that as we increase the scale of admixed genotype data with more complex structure, Mahalanobis distance is better suited as it is known to project correlated high dimensional data to an uncorrelated lower dimensional space where it recovers the hidden Euclidean distances [32].

**Fig. 8:**
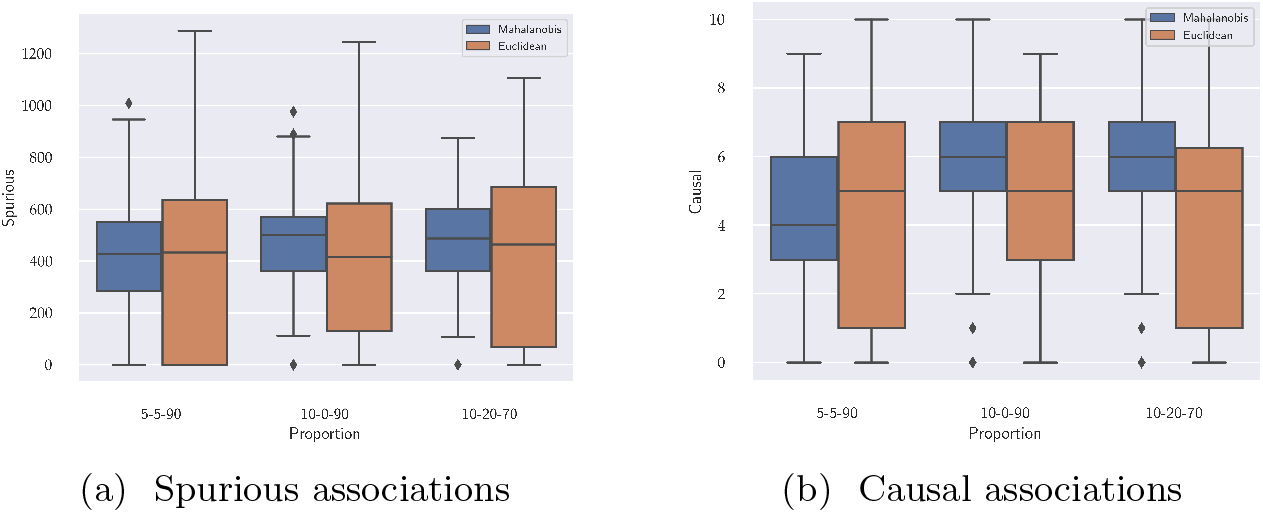
Box plots for spurious and causal associations obtained by running AHC with Mahalanobis and Euclidean distances on the PSD model (*α* = {0.1, 0.1, 0.1}). We observe similar performance on both the distance metrics in terms of identifying true causal variants. Mahalanobis distance discovers less spurious associations than Euclidean distance.

