## Supplementary Material for "CluStrat: a structure informed clustering strategy for population stratification"

### Appendix A

##### Data simulator

The complete simulation study on quantitative traits with population structure latent variable is constructed in 5 different ways for 3 different proportions of variance among genetic effects, non-genetic effects and random noise, all of which contributing to the trait. We simulated 100 independent datasets containing $m = 1,000$ individuals and $n = 100,000$ markers from a quantitative trait model 9. Let $Z$ be a latent variable which captures environmental factors contributed by population structure. Equation 9 allows interdependence of structure, lifestyle and environment. We assume $\mathbf{E}\left[\epsilon_j | z_j\right] \sim \mathcal{N}(0, \sigma^2(z_j))$ allowing for heteroskedasticity of the random noise variation [41]. Therefore, $x^j = (x_{1j}, x_{2j}, \cdots, x_{mj})^\top$, $\lambda_j$ and $\sigma^2$ can be thought of as functions of $z_j$ where $Z = (z_1, z_2, \cdots, z_m)$. $\lambda_j$ is unspecified but along with $z_j$, they are assumed to be dependent, random variables. Thus, the population genetic model is dependent on the structure variable $z_j$ for each individual. We define the corresponding binary trait model as

$$\log\left(\frac{\Pr\left(y_j = 1\right)}{\Pr\left(y_j = 0\right)}\right) = \alpha + \sum_{i=1}^{m} \beta_i x_{ij} + \lambda_j \tag{10}$$

using the Odds Ratio (OR) as the classifier for disease status from the continuous variable $y$.

The complete simulation study on quantitative traits with population structure latent variable is constructed in 5 different ways for 3 different proportions of variance among genetic effects, non-genetic environmental effects and random noise, all of which contributing to the trait. Therefore $\mathbf{Var}\left[\sum_{i=1}^{n} \beta_i x_{ij}\right]$, $\mathbf{Var}\left[\sum_{j=1}^{n} \lambda_j\right]$ and $\mathbf{Var}\left[\epsilon_j\right]$ are assigned in proportions of (5%,5%,90%), (10%,0%,90%) and (10%,20%,70%), respectively. Thus, we varied the amount of genetic contribution to the trait for each simulation scenarios and capture variable amounts of population structure confounding. We simulated ten truly associated SNPs whose effect sizes were distributed according to a Normal distribution and we set $\beta_i = 0$ for all other non-causal SNPs.

The genotype matrix $\mathbf{X} \in \mathbb{R}^{m \times n}$ consisting of the simulated allele frequencies was simulated using the algorithm from a previous study [25,41]. Specifically, we set $\mathbf{F} = \mathbf{TS}$ where $\mathbf{T} \in \mathbb{R}^{m \times d}$ and $\mathbf{S} \in \mathbb{R}^{d \times n}$ where $d \leq n$ is the number of population groups. $\mathbf{S}$ is the matrix containing the population groups encompassing the structure for the individuals shared across all SNPs. On the other hand, $\mathbf{T}$ characterizes how the structure is manifested in the allele frequencies of each SNP [25]. Finally, projecting $\mathbf{S}$ onto the column space of $\mathbf{T}$ we obtain the allele frequency matrix $\mathbf{F}$. We sample $\mathbf{X}$ as a special case of $\mathbf{F}$ for Balding-Nichols (BN), Pritchard-Stephens-Donelly (PSD) and TGP (1000 Genomes Project), respectively. We formed $\mathbf{T}$ and $\mathbf{S}$ for the above 5 simulations with 3 scenarios each and continuous traits, resulting in, 15 different evaluation scenarios each for continuous and binary traits. The algorithm for constructing $\mathbf{T}$ and $\mathbf{S}$ is detailed in reference [25,41].

For BN, the allele frequency matrix is simulated from the HapMap phase 3 dataset [14] using three unrelated populations. The final genotype matrix, $\mathbf{X}$, is drawn independently at random from the Binomial distribution with parameters $n$ set to 2, denoting the allele status (0,1 or 2) corresponding to homozygous major/minor or heterozygous with probability $p$ set to the simulated allele frequency for each individual-SNP pair. For PSD, the allele frequency matrix was drawn from the BN frequency distribution. However, it differs from BN in simulating $\mathbf{S}$ by i.i.d draws from Dirichlet distribution with varying $\alpha$ which denotes the parameter influencing the relatedness between the individuals. We show results for $\alpha = \{0.01, 0.1, 0.5\}$ here and conducted simulations on a wide range of $\alpha$ values from 0.01 to 0.5.

##### Real dataset

To capture real world population structure, we applied CluStrat on the Parkinson's Disease (PD) data available from the The Wellcome Trust Case Control Consortium (WTCCC2) study containing 4706 individuals (2837 controls and 1869 cases) across 517,672 SNPs. After performing quality control by filtering for genotyping rate lower than 99%, MAF less than 0.01 and Hardy-Weinberg equilibrium less than 0.001 and pruning for LD between variants higher than 0.2 squared correlation we obtained 99,631 markers.

##### Distance metrics for Hierarchical clustering

CluStrat computes the distance matrix $\mathbf{D}$ from $\mathbf{X}$ to perform the AHC. The choice of distance metric is user defined. However, we choose the distance metric based on LD induced distances to capture the cryptic relatedness between individuals in a population which is not otherwise captured by other stratification methods. We use the normalized genotype matrix $\mathbf{X}$ following the standard normalization procedure by minor allele frequency of each marker. Let us consider the unscaled GRM which captures the Euclidean distances as $\mathbf{D} = \mathbf{X}\mathbf{X}^\top$ and let $\mathbf{I} \in \mathbb{R}^{n \times n}$ (in the order of the number of markers $n$). Thus $\mathbf{D}$ can be rewritten as

$$\mathbf{D} = \mathbf{X}\mathbf{I}\mathbf{X}^\top \tag{11}$$

Thus we can see the unscaled GRM as the same weighting on the diagonal for all markers. In an arbitrarily structured breeding population, there exists correlation between loci due to linkage resulting in varying values along the diagonal or a block-diagonal structure in the GRM. Thus, it is important to account for this LD covariance structure in the computation of the GRM [34]. One way to account for the LD structure in GRM is to use the squared Mahalanobis distance [32,36] (denoted as $\mathbf{D}$ for simplification). Given a matrix $\mathbf{G} \in \mathbb{R}^{n \times n}$ which contains the covariance structure of LD (covariance due to markers), then the

LD-corrected GRM with Mahalanobis distance is defined as

$$\mathbf{D} = \mathbf{X}\mathbf{G}^{-1}\mathbf{X}^\top \tag{12}$$

The RHS of equation 1 represents the squared multivariate Mahalanobis distance between individuals. Mahalanobis distance is useful in a high-dimensional setting where the Euclidean distances fail to capture the true distances between observations. It achieves this by taking correlation between the features captured in the SNP covariance matrix into account. The Cholesky factorization of the covariance matrix $\mathbf{G} = \mathbf{L}\mathbf{L}^\top$ where $\mathbf{L}$ is the lower diagonal matrix known as the Cholesky factor of $\mathbf{G}$ [34]. We can represent equation 1 as

$$\begin{aligned}
\mathbf{X}\mathbf{G}^{-1}\mathbf{X}^\top &= \mathbf{X}\left(\mathbf{L}\mathbf{L}^\top\right)^{-1}\mathbf{X}^\top \\
&= \mathbf{X}(\mathbf{L}^\top)^{-1}(\mathbf{L})^{-1}\mathbf{X}^\top \\
&= \left(\mathbf{X}(\mathbf{L}^{-1})^\top\right)\left(\mathbf{L}^{-1}\mathbf{X}\right)^\top \\
&= \left(\mathbf{L}^{-1}\mathbf{X}^\top\right)^\top\left(\mathbf{L}^{-1}\mathbf{X}^\top\right) \\
&= \mathbf{Q}^\top\mathbf{Q}
\end{aligned}$$

$\mathbf{Q} = \mathbf{L}^{-1}\mathbf{X}^\top$ represents the transformed variables and $\mathbf{Q}^\top\mathbf{Q}$ is the squared Euclidean distance between the transformed variables. Thus, Mahalanobis distance accounts for covariance between variables by transforming the data into an uncorrelated form and computing the euclidean distances between them.

**Mahalanobis Distance and Leverage Scores** Mahalanobis distance is known to be connected to statistical leverage [47], which is extended in the RandNLA framework as leverage scores. We show this relationship by first noting that Mahalanobis distance is invariant to linear transformations, which means the Mahalanobis distance between two vectors,

$$\mathbf{D}(\mathbf{X}_{i*}, \mathbf{X}_{j*}) = (\mathbf{X}_{i*} - \mathbf{X}_{j*})\mathbf{G}^{-1}(\mathbf{X}_{i*} - \mathbf{X}_{j*})^\top \tag{13}$$

can have zero means for each vector. In our genotype matrix, $\mathbf{X} \in \mathbb{R}^{m\times n}$, we have $n$ markers and $m$ observations. The design matrix $\mathbf{X}$ on which we intend to fit the model, however, must contain an intercept and thus we refer to $\mathbf{X}$ here as the design matrix containing the intercept column followed by one column for each SNP for all the individuals in rows. Furthermore, as we compute the Mahalanobis distance with respect to the *low-rank* genotype matrix $\mathbf{X}_k$, we only consider the *low-rank leverage scores* (rather than the leverage scores of the original matrix $\mathbf{X}$) which are essentially the diagonal elements of the following projection-matrix:

$$\mathbf{H} = \mathbf{X}_k\left(\mathbf{X}_k^\top\mathbf{X}_k\right)^{-1}\mathbf{X}_k^\top \tag{14}$$

and similarly, the off-diagonal elements of $\mathbf{H}$ are called *cross-leverage scores* of $\mathbf{X}_k$. Now, we will give a clean connection between Mahalanobis distance and these leverage and cross-leverage scores.

First, consider the diagonal elements of $\mathbf{H}$ *i.e.* when $i = j$, we have

$$\mathbf{H}_{ii} = (1; \mathbf{X}_{k_{i*}}) \left(\mathbf{X}_k^\top \mathbf{X}_k\right)^{-1} (1; \mathbf{X}_{k_{i*}})^\top . \tag{15}$$

Exploiting the structure of $\left(\mathbf{X}_k^\top \mathbf{X}_k\right)^{-1}$, we can reformulate it in terms of a block matrix as follows

$$\mathbf{X}_k^\top \mathbf{X}_k = m \begin{pmatrix} 1 & \mathbf{0}^\top \\ \mathbf{0} & \mathbf{C} \end{pmatrix}$$

where $\mathbf{C}_{ij} = \frac{1}{m} \sum_{\ell=1}^m \mathbf{X}_{k_{\ell i}} \mathbf{X}_{k_{\ell j}} = \frac{m-1}{m} Cov(\mathbf{X}_{k_{*i}}, \mathbf{X}_{k_{*j}}) = \frac{m-1}{m} \mathbf{\Sigma}_{ij}$. $\mathbf{\Sigma}$ here is the corresponding sample covariance matrix. Thus,

$$\left(\mathbf{X}_k^\top \mathbf{X}_k\right)^{-1} = \frac{1}{n} \begin{pmatrix} 1 & \mathbf{0}^\top \\ \mathbf{0} & \mathbf{C}^{-1} \end{pmatrix} = \begin{pmatrix} \frac{1}{n} & \mathbf{0}^\top \\ \mathbf{0} & \frac{1}{n-1} \mathbf{\Sigma^{-1}} \end{pmatrix}$$

From Equation 15 we obtain

$$\mathbf{H}_i = (1; \mathbf{X}_{k_{i*}}) \begin{pmatrix} \frac{1}{m} & \mathbf{0}^\top \\ \mathbf{0} & \frac{1}{m-1} \mathbf{\Sigma^{-1}} \end{pmatrix} (1; \mathbf{X}_{k_{i*}})^\top \tag{16}$$

$$= \frac{1}{m} + \frac{1}{m-1} \mathbf{X}_{k_{i*}} \mathbf{\Sigma^{-1}} \mathbf{X}_{k_{i*}}^\top \tag{17}$$

$$= \frac{1}{m} + \frac{1}{m-1} \mathbf{D}\left(\mathbf{X}_{k_{i*}}, 0\right) \tag{18}$$

Solving for

$$\mathbf{D}_i = \mathbf{D}(\mathbf{X}_{k_{i*}}, 0)$$

yields,

$$\mathbf{D}_i = (m-1) \left(\mathbf{H}_i - \frac{1}{m}\right)$$

Similarly, we can prove the cross-leverage scores

$$\mathbf{H}_{ij} = \frac{1}{m} + \frac{1}{m-1} \mathbf{X}_{k_{i*}} \mathbf{\Sigma^{-1}} \mathbf{X}_{k_{j*}} \tag{19}$$

To prove the relationship of $\mathbf{H}_{ij}$ with $\mathbf{D}_{ij}$ we see,

$$\begin{aligned} \mathbf{D}(\mathbf{X}_{k_{i*}}, \mathbf{X}_{k_{j*}}) &= (\mathbf{X}_{k_{i*}} - \mathbf{X}_{k_{j*}}^\top) \mathbf{\Sigma}^{-1} (\mathbf{X}_{k_{i*}} - \mathbf{X}_{k_{j*}}) \\ &= \mathbf{D}(\mathbf{X}_{k_{i*}}, 0) + \mathbf{D}(\mathbf{X}_{k_{j*}}, 0) - 2\mathbf{X}_{k_{i*}} \mathbf{\Sigma}^{-1} \mathbf{X}_{k_{j*}} \\ &= (m-1)(\mathbf{H}_i - \frac{1}{m}) + (m-1)(\mathbf{H}_j - \frac{1}{m}) - 2(m-1)(\mathbf{H}_{ij} - \frac{1}{m}) \\ &= (m-1)(\mathbf{H}_i + \mathbf{H}_j - 2\mathbf{H}_{ij}) \end{aligned}$$

If we take $\mathbf{X}_{k_{i*}} = \mathbf{X}_{k_{j*}}$ then we find $\mathbf{D}(\mathbf{X}_{k_{i*}}, \mathbf{X}_{k_{j*}}) = 0$. Thus, we show that Mahalanobis distance between two vectors can be computed by the corresponding vector's leverage scores.

One of the key computational bottlenecks of Mahalanobis distance is computing the inverse of the SNP covariance matrix $\mathbf{G}$ as required in Equation 1. In real datasets, with the improvements in genotyping and sequencing technologies, the number of SNPs can be in the millions, thereby making $\mathbf{G}$ in the order of million times million and infeasible to store in secondary memory. Here, we propose the first approximation of Mahalanobis distance by computing leverage and cross-leverage scores in a faster and efficient way. As we have shown in Equation 19 and 16 following up from previous work [47], Mahalanobis distance can be written in terms of leverage scores. Advances in RandNLA community have brought about faster computations for leverage scores as well as cross-leverage scores; hence, we can compute approximations to these scores using random sampling algorithms with theoretical guarantees [19]. For our purposes of demonstrating the proof-of-concept, we work with simulated data as described above for 1,000 individuals and 500,000 SNPs which could be feasibly processed in a personal workstation to compute the deterministic leverage and cross-leverage scores.

**Computing leverage and cross-leverage scores.** In fact, we do not need to compute the *rank-k* leverage and cross-leverage scores exactly. Using the idea of [19,12], they can be well-approximated in a much faster way with high-probability. In particular, computing $m$ row-leverage scores takes time

$$\mathcal{O}\left(\mathrm{nnz}(\mathbf{X}_k)\log n + k^3 \log^2 k + k^2 \log n\right) ,$$

where nnz means the non-zero entries of the matrix, and computation of the high-valued cross-leverage scores can be done in time

$$\mathcal{O}(\mathrm{nnz}(\mathbf{X}_k)\log^3 n) .$$

##### Fast Computation of Standard errors

For biobank-scale data-sets requiring terabytes of memory, computing the standard error can be a challenge. However, we can use random projection based sketching matrices to find an approximate standard error for each marker by projecting the genotype matrix $\mathbf{X}$ on a sketching matrix $\mathbf{S} \in \mathbb{R}^{n \times r}$ to form a sketch $\mathbf{XS}$. We can rewrite the standard error in Equation 3 to find it's approximate as,

$$\tilde{\hat{SE}}(\hat{\beta}_i) = \sigma^2 \| \left(\mathbf{XSS}^\top \mathbf{X}^\top + \lambda \mathbf{I}_m\right)^{-1} \mathbf{X}^{(i)} \|_2^2 \quad (20)$$

The sketched matrix $\mathbf{XS}$ generically has the same rank but much fewer columns than $\mathbf{X}$, satisfying $1 \leq r \leq min\{m, n\}$. Sketching, in general, is used to speed up solving systems of linear equations [20,22,12]. The sketching dimension, $r$, is directly proportional to the accuracy obtained by the approximate standard errors. Some prior knowledge of the design matrix, $\mathbf{X}$, helps determine the target rank, $r$, that will result in satisfactory error guarantees. The sketching matrix, $\mathbf{S}$, can be chosen simply as i.i.d normal random variables with mean equal to zero and variance equal to $\frac{1}{r}$. There exists other ways to choose $\mathbf{S}$ based on random projections as shown in previous work involving Fast Johnson-Lindenstrauss Transform [1], Subsampled Randomized Hadamard Transform [20,21] and Count-Sketch matrices [9] from streaming setting involving faster computation with sparse matrices.

**Time to compute eqn.** (20)**.** Following the discussion as in [11], let the time to compute the sketch $\mathbf{XS} \in \mathbb{R}^{m \times s}$ be $T(\mathbf{X}, \mathbf{S})$ which depends on the particular construction of $\mathbf{S}$. In order to invert the matrix $\mathbf{Q} = \mathbf{XSS}^\top\mathbf{X}^\top$, it suffices to compute the SVD of the matrix $\mathbf{XS}$. Notice that given the singular values of $\mathbf{XS}$, we can compute the singular values of $\mathbf{Q}$ and also notice that the left and right singular vectors of $\mathbf{Q}$ are the same as the left singular vectors of $\mathbf{XS}$. Interestingly, we do not need to compute $\mathbf{Q}^{-1}$. Instead, we can store it implicitly by storing the left (and right) singular vectors of $\mathbf{Q}$ along with its singular values, $\mathbf{\Sigma_Q}$. Then, we can compute all necessary matrix-vector products using this implicit representation of $\mathbf{Q}^{-1}$. Thus, inverting $\mathbf{Q}$ takes $\mathcal{O}\left(sm^2\right)$ time and this will eventually dominate the computation of all other matrix-vector products and the Euclidean-norm. Therefore, total running time to compute eqn. (20) is given by

$$\mathcal{T} = \mathcal{O}(sm^2) + T(\mathbf{X}, \mathbf{S})$$

Clearly, specific constructions of the sketching matrix $\mathbf{S}$ will determine both $s$ and $T(\mathbf{X}, \mathbf{S})$, and therefore $\mathcal{T}$. For example, if $\mathbf{S}$ is a subsampled randomized Hadamard transform (SRHT) matrix, then we have, $T(\mathbf{X}, \mathbf{S}) = \mathcal{O}(mn \log n)$ and $s = \Omega((m \log(m/\delta))/\varepsilon^2)$; therefore $\mathcal{T} = \mathcal{O}((m^3 \log(m/\delta))/\varepsilon^2) + \mathcal{O}(mn \log n)$. Similarly, if $\mathbf{S}$ has sub-Gaussian entries, then $s = \mathcal{O}(m/\varepsilon^2)$ and $T(\mathbf{X}, \mathbf{S}) = \mathcal{O}(m^2 n)$; therefore $\mathcal{T} = \mathcal{O}(m^2 n)$. Furthermore, if $\mathbf{S}$ is a count-sketch matrix of [12], then, in this case, $s = \Omega(\frac{m^2}{\varepsilon^2 \delta})$ and $T(\mathbf{X}, \mathbf{S}) = \mathrm{nnz}(\mathbf{X})$. So, total running time $\mathcal{T} = \mathcal{O}(\mathrm{nnz}(\mathbf{X}) + \frac{m^4}{\varepsilon^2 \delta})$. Here, $\varepsilon$ is the accuracy parameter and $\delta$ is the corresponding failure probability.

Note that sketching-dimension, $s$, for sub-Gaussian is optimal, but $T(\mathbf{X}, \mathbf{S})$ takes much time. On the other hand, for count-sketch, $T(\mathbf{X}, \mathbf{S})$ is much faster (only $\mathrm{nnz}(\mathbf{X})$), but sketching-dimension is huge $\mathcal{O}(m^2/\varepsilon^2)$. In a recent work [13], the authors showed that we can actually use all the sketches discussed here in conjunction with each other to get the best performance both in terms of sketching-dimension as well as computation time. More precisely, if one set $\mathbf{S} = \mathbf{S}_1\mathbf{S}_2\mathbf{S}_3$ with $\mathbf{S}_1$ being the count-sketch, $\mathbf{S}_2$ being the SRHT and $\mathbf{S}_3$ being the sub-Gaussian , then $\mathbf{XS}$ will have $\mathcal{O}(m/\varepsilon^2)$ columns with running time $T(\mathbf{X}, \mathbf{S}) = \mathcal{O}(\mathrm{nnz}(A)) + \tilde{\mathcal{O}}\left(\varepsilon^{-\mathcal{O}(1)}\left(m^3 + m^2 d\right)\right)$.

### Appendix B

##### Comparing Stratification Methods

**BN model** The BN model simulates scenarios with unrelated isolated populations (Figure 2 (i)) and serves as the basic case for arbitrarily structured population with no admixture.

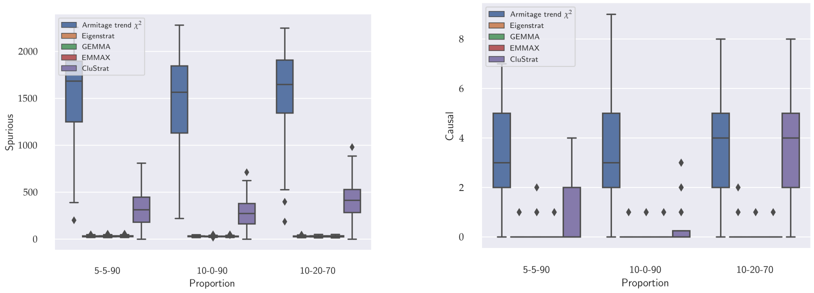


(a) Spurious associations (b) Causal associations

Fig. 4: Box plots for spurious and causal associations on the BN model shows that Armitage trend $\chi^2$ has the maximum number of spurious associations containing about 4-5 causal SNPs whereas EIGENSTRAT has minimum number of spurious associations while detecting almost zero causal SNPs. CluStrat has more spurious associations than EIGENSTRAT and considerably less than Armitage trend $\chi^2$ recovering slightly more number of causal SNPs than the latter.

The samples when projected on the top two PCs clearly resembles three isolated clusters with no connections between them. This is an ideal case when the populations are not mixing due to environmental factors acting as barriers of gene flow between populations. GWAS has shown to be robust in these settings [45]; however, the cryptic relatedness for each cluster remains a plaguing issue [5]. We ran CluStrat on this scenario with p-value threshold set to $p = \frac{25}{m_i} = 0.0025$ ($m_i$ is the number of SNPs in each iteration, set to 10,000 for 100 iterations). The expected number of spurious association as mentioned in [41] is $m_0 \times p$ where $m_0 = m-$ number of causal SNPs. In our case, as we set the number of causal SNPs to 10 as per [41], $m_0 = 9990$ and therefore, the number of spurious associations to be approximately 25 with degree of freedom set to 1 for genotypes.

Armitage trend $\chi^2$ with no population structure correction renders almost half of the SNPs in the simulation study as true associations resulting in a considerable amount of spurious associations highlighting the need for population structure correction. EIGENSTRAT on the other hand results in the expected number of spurious associations as also shown in previous work [38]. However, it behaves stringently and detects zero causal SNPs almost all of the time (Figure 4). CluStrat, however, strikes a balance between the two and generates far more spurious associations than the expected value but about 5 folds less than Armitage trend $\chi^2$ recovering a slightly higher number of causal SNPs. This shows that in the ideal case of population structure correction, CluStrat can identify more causal SNPs due to the structure informed clustering setup which widely used stratification correction methods lack.

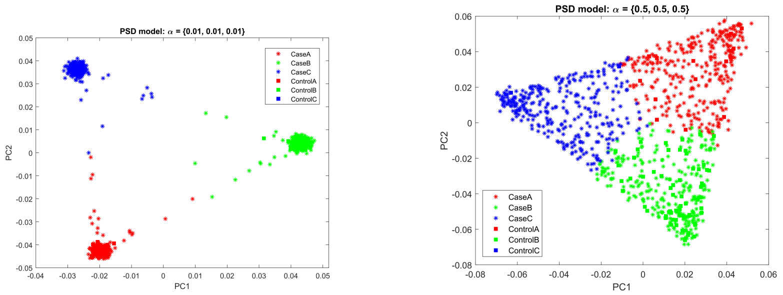


(a) $\alpha = \{0.01, 0.01, 0.01\}$ (b) $\alpha = \{0.5, 0.5, 0.5\}$

Fig. 5: Projection of the samples from PSD model with varying sets of values of $\alpha$. We observe that increasing $\alpha$ increases the density between individuals leading to admixture and creates a uniform gradient as all values of $\alpha_i$ are equal.

**PSD model** The PSD model emulates real world datasets more closely than BN model. It allows for admixing individuals and gradients across the populations. It is sampled from the Dirichlet distribution parameterized by a concentration parameter $\alpha \in \mathbb{R}^d$ where $d = 3$ (the number of populations for all simulations conducted). A higher value of $\alpha_i$ corresponds to greater weight of $i^{th}$ population. We ran CluStrat on the PSD model with varying number of $\alpha$ from 0.01 to 0.5 and kept equal $\alpha_i$ for a symmetric distribution. We report the boxplots of spurious and causal associations (Figure 6 and 7) for $\alpha = 0.1, 0.5$ and and observe that for the first case of variance, $(5\%, 5\%, 90\%)$, Armitage trend $\chi^2$ and CluStrat performs almost similarly in terms of spurious associations. This is due to the fact that only 5% of the trait is explained by true genetic associations in presence of LD and the rest is noise and environmental factors. However, CluStrat outnumbers EIGENSTRAT, GEMMA and EMMAX in terms of causal associations and detects four to six fold more true causal SNPs. For the other two variance proportions, CluStrat performed better than the other methods in detecting the causal associations and strikes a balance in terms of spurious associations.

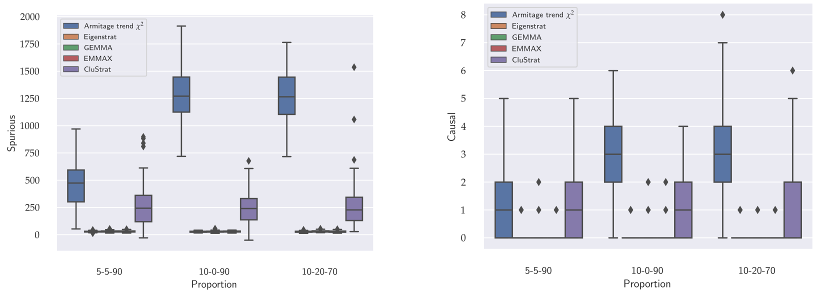


(a) Spurious associations (b) Causal associations

Fig. 6: Box plots for spurious and causal associations on the PSD model ($\alpha = \{0.1, 0.1, 0.1\}$) shows Armitage trend $\chi^2$ has maximum number of spurious associations containing less causal SNPs than the BN model (Figure 4) owing to the admixed nature of the individuals in PSD. EIGENSTRAT, GEMMA and EMMAX has least number of spurious associations while detecting almost zero causal SNPs. CluStrat has more spurious associations than the standard approaches and less than Armitage trend $\chi^2$ while recovering two to three fold more causal SNPs.

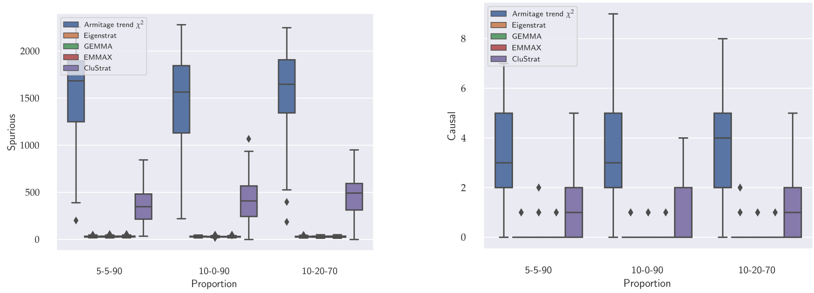


(a) Spurious associations (b) Causal associations

Fig. 7: Box plots for spurious and causal associations on the PSD model ($\alpha = \{0.5, 0.5, 0.5\}$) shows Armitage trend $\chi^2$ has maximum number of spurious associations containing less causal SNPs than the BN model (Figure 4) owing to the overtly admixed nature of the individuals in PSD. EIGENSTRAT, GEMMA and EMMAX has least number of spurious associations while detecting almost zero causal SNPs. CluStrat has more spurious associations than the standard approaches and slightly more than $\alpha = 0.1$ owing to more admixed nature of the data. It has considerably less spurious associations than Armitage trend $\chi^2$ while recovering two to three fold more causal SNPs.

#### Comparing Distance metrics

CluStrat with Euclidean distance metric based GRM (sample covariance matrix) also contains structure information as part of the relationships between the individuals within and between population groups. The GRM with Euclidean distance is straightforward to compute as shown below

$$\mathbf{D} = \mathbf{X}^\top \mathbf{X}$$

where $\mathbf{X} \in \mathbb{R}^{n \times m}$, with number of markers, $n$ and number of samples, $m$ ($n >> m$). We show that although Euclidean distances between individuals is straightforward to compute, it fails to distinguish fine-grained distances between individuals in the same cluster owing to cryptic relatedness. This is highlighted after performing AHC using Ward's linkage method which minimizes the increase in sum of squares between two cluster centroids in order to decide when to merge them. (Figure 1).

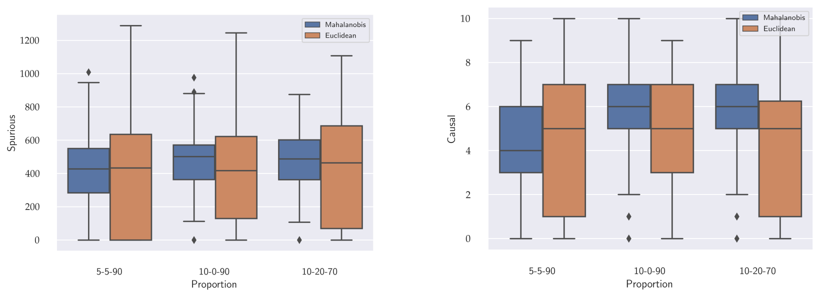


(a) Spurious associations (b) Causal associations

Fig. 8: Box plots for spurious and causal associations obtained by running AHC with Mahalanobis and Euclidean distances on the PSD model ($\alpha = \{0.1, 0.1, 0.1\}$). We observe similar performance on both the distance metrics in terms of identifying true causal variants. Mahalanobis distance discovers less spurious associations than Euclidean distance.

When Mahalanobis distance based GRM is used instead of Euclidean distance in AHC on PSD model with 1,000 individuals and 10,000 SNPs across 3 admixed arbitrarily structured ethnic groups, it reveals four broad clusters with various fine-grained sub-clusters revealing how Mahalanobis distance help recover cryptic relatedness and substructure within a population.

Due to admixture in the PSD model ($\alpha = \{0.1, 0.1, 0.1\}$) as shown in Figure 5 the dendrogram finds three broad clusters owing to the three populations in the simulation. It subsequently finds different sub-clusters at different depth on the horizontal axis. Thus, identifying interaction between individuals inside a cluster. This is a significant advantage of using Mahalanobis distance over it's Euclidean counterpart as the latter only reveals three broad clusters with indistinguishable interactions in each cluster (Figure 1).

When we ran AHC with both the distances, we observe similar performance on the PSD model with Mahalanobis distance based GRM performing slightly better with respect to it's Euclidean counterpart (Figure 8). We note that as we increase the scale of admixed genotype data with more complex structure, Mahalanobis distance is better suited as it is known to project correlated high dimensional data to an uncorrelated lower dimensional space where it recovers the hidden Euclidean distances [32].

Table 1: Table showing strongest associations after running CluStrat on WTCCC2 PD data

| Chrom# | SNPID | GeneID | p-value |
|---|---|---|---|
| 2 | rs10177996 | WNT10A | $2.22 \times 10^{-16}$ |
| 2 | rs1059823 | SLC11A1 | $2.4 \times 10^{-16}$ |
| 4 | rs11936554 | UNC5C | $4 \times 10^{-16}$ |
| 2 | rs13013415 | WDR33 | $4.5 \times 10^{-16}$ |
| 2 | rs1509467 | MAP2 | $5 \times 10^{-15}$ |
| 3 | rs1516570 | GRM7 | $5.7 \times 10^{-14}$ |
| 6 | rs176713 | BACH2 | $6 \times 10^{-12}$ |
| 6 | rs176713 | BACH2 | $6 \times 10^{-12}$ |
| 4 | rs2322559 | SLIT2 | $6.1 \times 10^{-12}$ |
| 4 | rs2328457 | AIG1 | $6.13 \times 10^{-12}$ |
| 3 | rs3816969 | NMNAT3 | $6.15 \times 10^{-12}$ |
| 3 | rs4677964 | PARP15 | $7.15 \times 10^{-10}$ |
